# AN INTEGRATED HEART-ON-A-CHIP NAM RESOLVES DISTINCT CLINICAL PHENOTYPES OF MYOCARDIAL ISCHEMIA-REPERFUSION INJURY

**DOI:** 10.64898/2026.10.23.761466

**Authors:** Khanh Tran, Ashley Roselynn Vincent, Caroline Whitmer, Rami N. Khayat, Anna Grosberg

## Abstract

**Background:** Reperfusion is required for myocardial salvage yet itself injures the heart, and cardioprotective interventions that reduce infarct size experimentally have repeatedly failed to improve outcomes in patients. These interventions are evaluated against viability or infarct size alone, whereas ischemia-reperfusion (I/R) injury produces structural, biochemical, and contractile dysfunction in the same tissue. No *in vitro* new approach methodology (NAM), however, resolves all three within one system.

**Methods:** A NAM was created within a heart-on-a-chip platform, which was constructed by culturing neonatal rat ventricular myocytes (NRVMs) on muscular thin films. Cardiomyocytes were subjected to normoxia, 2 hours of ischemia, and ischemia followed by 2 hours of reperfusion. Sarcomeric architecture was quantified by immunofluorescence, cardiomyocyte injury by LIVE/DEAD staining and cardiac troponin I release into conditioned media, while contractile stress and spontaneous beat frequency were recorded on the same films before and after each condition.

**Results:** Under ischemia, z-line density decreased while architectural metrics were preserved, indicating cell loss rather than sarcomere disorganization. Furthermore, viability decreased, troponin elevated, and every contractile stress component declined, while spontaneous beat frequency was unchanged. Reperfusion fractured this uniform depression. Cell death and troponin release increased further, yet structure was unchanged from ischemia, so every functional change occurred on a depleted but architecturally intact cardiomyocyte population. Systolic stress recovered partially, whereas diastolic stress exceeded baseline, leaving active stress suppressed despite oxygen recovery. Beat frequency increased with pronounced variability.

**Conclusions:** This heart-on-a-chip NAM resolved structural, biochemical, and contractile injury that global indices conflate, distinguishing ischemic from reperfusion pathophysiology. Interestingly, individual readouts corresponded to distinct clinical syndromes of ischemic injury, positioning this platform for future testing of cardioprotective candidates against the full injury phenotype.

## 1. INTRODUCTION

Modern interventional cardiology has transformed the prognosis of acute myocardial infarction (MI). Indeed, door-to-balloon times have fallen, stent technology has advanced, and antithrombotic regimens have been refined^1–3^, yet mortality has plateaued and the number of patients developing post-myocardial infarction (MI) heart failure continues to rise^4–7^. A central contributor to this persistent burden is myocardial ischemia-reperfusion (I/R) injury^8^. Paradoxically, the restoration of coronary blood flow — while essential for myocardial salvage — triggers a cascade of injurious events, including mitochondrial permeability transition pore (mPTP) opening, intracellular calcium overload, and reactive oxygen species generation, that collectively accelerate cardiomyocyte death and impair contractile recovery^8–11^. Reperfusion injury has been estimated to account for up to 50% of the final infarct size^8,9^, and its clinical footprint is substantial; cardiac MRI studies identify microvascular obstruction in 40–50% of reperfused ST-elevation myocardial infarction (STEMI) patients^12,13^. Despite decades of investigation, the numerous cardioprotective strategies that have demonstrated promising infarct size reduction in experimental studies have consistently failed to improve outcomes in patients^14–18^. One underlying cause may be that candidate interventions are evaluated against narrow, single-dimension endpoints, most commonly infarct size or cell viability, while I/R injury manifests clinically as simultaneous structural, biochemical, and contractile dysfunction^19^. New approach methodologies (NAMs) could address this limitation by allowing structural, biochemical, and contractile injury to be quantified within an *in vitro* system.

Isolated cardiomyocyte hypoxia-reoxygenation systems offer precise control of the cellular and extracellular environment but rely predominantly on viability assays as their primary endpoint^20^, providing no measure of contractile function or sarcomeric integrity. Three-dimensional organoid and microtissue models improve structural organization but remain similarly narrow. Organoids characterize injury largely through cell death^21,22^, whereas spheroid models add sarcomere disruption and core apoptosis without functional measurement^23^, and both lack the electromechanical conditioning and real-time force measurement required to track functional recovery^24^. Other chip-based systems infer injury indirectly, through beating variability and lactate accumulation^25^ or extracellular vesicle-associated miRNA release^26^, without capturing structural information. Engineered heart tissue and other cantilever-based platforms pair viability or biochemical markers with contractile force measurement, but neither resolves sarcomeric architecture at cellular resolution^27,28^. These efforts complement the multicellular three-dimensional and organ-on-chip approaches that have advanced cardiac injury modeling in recent years^29,30^. Consequently, resolving the clinical I/R phenotype *in vitro* requires a platform that measures structural, biochemical and contractile injury within the same system, to evaluate candidate cardioprotective interventions against the full clinical injury phenotype.

Here, we present an integrated *in vitro* platform for the study of myocardial ischemia and I/R injury, characterized across structural, biochemical, and contractile axes in the same system. We demonstrate that each readout of this NAM captures a distinct and clinically recognizable dimension of the I/R injury phenotype, and that together they recapitulate the multidimensional injury continuum observed as a result of ischemia and reperfusion.

## 2. METHODS

### Muscular thin film (MTF) fabrication

MTFs were fabricated following an established protocol^31–34^. Briefly, large glass coverslips (76 × 83 mm^2^; Brain Research Laboratories, Newton, MA) were sonicated in ethanol for 10 min and allowed to air dry. Protective film strips were placed parallel to the coverslip edges approximately 1 cm apart to mask discrete regions of the glass surface. Poly(N-isopropylacrylamide) (PIPAAm; Sigma-Aldrich, Burlington, MA) was spin-coated onto the unmasked glass surface and allowed to cure at room temperature for at least 10 min. After removing the protective strips, polydimethylsiloxane (PDMS; Sylgard 184, Ellsworth Adhesives, Germantown, WI) was prepared at a 10:1 base-to-curing-agent ratio then spin-coated over the entire coverslip surface. These yielded alternating regions coated with PDMS alone (over the formerly masked areas) and with PIPAAm beneath the PDMS layer (over the formerly unmasked areas). Substrates were then fully cured overnight at 65°C. The cured PDMS was lightly scored to define four discrete thin-film strips without engraving the underlying glass, and individual chips (∼12.7 × 13.8 mm²) were separated using a laser cutter (Trotec Speedy 360, Trotec Laser GmbH, Plymouth, MA). Three chips from each large coverslip were reserved for thickness verification; PDMS film thicknesses were confirmed to fall within 12–15 µm by profilometry (Dektak 6M, Veeco Instruments, Plainview, NY). Fibronectin (FN; Sigma-Aldrich, St. Louis, MO) was patterned onto the PDMS surface by microcontact printing^31,33^. PDMS stamps bearing parallel lines 22 μm wide with 3 μm gaps were sterilized in ethanol and dried under sterile conditions. The patterned stamp surface was inked with 50 μg/mL of FN for 1 h. Coverslip surfaces were simultaneously sterilized and activated by UV-ozone treatment for 8 minutes (Jelight Company, Irvine, CA). Stamps were dried under compressed nitrogen and brought into conformal contact with the activated coverslips to transfer the FN pattern. Chips were then housed in 12-well plates pre-coated with 1% Phuronics F-127 (BASF Group, Parsippany, NJ) in deionized water for 10 minutes and subsequently washed three times with phosphate-buffered saline (PBS; ThermoFisher, Waltham, MA; Cat# 10010049). Chips were submerged in PBS and stored at 4°C until use, with all chips used within 4–7 days of FN stamping. The fibronectin micropattern guides unidirectional cardiomyocyte alignment, recapitulating the anisotropic architecture characteristic of native myocardial tissue^33,35,36^ (Fig. S1).

### Isolation and Culture of Neonatal Rat Ventricular Myocytes (NRVMs)

All animals for the study were treated according to the Institutional Animal Care and Use Committee of University of California, Irvine guidelines (IACUC Protocol No. 2025-020) and the NIH Guide for the Care and Use of Laboratory Animals. Ventricular myocardium was extracted from 2-day-old Sprague-Dawley rats (Charles River Laboratories, Wilmington, MA) as previously described^37–39^. Ventricular tissue was extracted from each pup, washed in Hanks’ balanced salt solution (HBSS; Thermo Fisher, Waltham, MA; Cat# 14170161), and subjected to overnight trypsin digestion (1 mg/mL in HBSS; Sigma-Aldrich, St. Louis, MO) for 12 h at 4°C on a rocker. Trypsin activity was neutralized with warm M199 culture media (Thermo Fisher, Waltham, MA; Cat# 11150067) supplemented with 10% fetal bovine serum (FBS; Avantor Seradigm, Select Grade, VWR, Radnor, PA; Cat# 76419-588) at 37°C. Then, the tissue was digested four times with 1 mg/mL collagenase type II (Worthington Biochemical Corporation, Lakewood, NJ) dissolved in HBSS at 37°C. The resulting cell suspension was washed in chilled HBSS and pelleted by centrifugation at 1,000 rpm for 7 min. Cells were resuspended in warm M199 media supplemented with 10% FBS and subjected to three consecutive 1-h differential pre-plating steps at 37°C to isolate the cardiomyocytes via the different adhesion rates of cardiomyocytes and fibroblasts. The purified cardiomyocyte fraction was collected and seeded onto fibronectin-patterned chips at a density of 2.7 × 10³ cells/mm² in 12-well plates. Cells were cultured in M199 media supplemented with 20 mM D-glucose, 10 mM HEPES, 2 mM L-glutamine, 0.1 mM minimum essential media (MEM) nonessential amino acids, 1.5 μM vitamin B-12, penicillin (50 U/mL), and 30% heat-inactivated fetal bovine serum (FBS) for 2 days. At 48 h, non-adherent and dead cells were removed by washing with PBS, and media was exchanged to 10% FBS M199 for the remainder of the culture period. All assays were performed 4 days after initial seeding (Fig. S1).

### Experimental Conditions

Six experimental conditions were compared (Fig. 3B): (1) Normoxia, in which MTFs were maintained under standard incubator conditions for 4 h as a control; (2) H_2_O_2_, in which cells were exposed to 100 μM hydrogen peroxide in M199 media for 4 h as a positive control for oxidative injury; (3) Ischemia only 2 h, in which cultures were subjected to 2 h of ischemia; and (4) Ischemia-Reperfusion 4 h (I/R), in which 2 h of simulated ischemia was followed by 2 h of reperfusion under normoxic conditions.

Two delayed control conditions were included to separate the effect of reperfusion from time-dependent changes in the preparation. NRVMs continue to mature in culture, so a change in contractile stress between 0 h baseline and a 4 h endpoint could reflect elapsed culture time rather than the experimental stimulus. In the delayed normoxia and delayed ischemia conditions, MTFs were maintained under standard incubator conditions for an initial 2 h without measurement (t_0_ to t_2_). Baseline contractility was recorded at t_2_, the time point at which reperfusion begins in the I/R condition. Delayed normoxia MTFs were then maintained under normoxia for a further 2 h, whereas delayed ischemia MTFs were subjected to 2 h of simulated ischemia. Endpoint contractility was recorded at t₄ in both conditions. Structural, viability, and cardiac troponin I (cTnI) assays were performed on the Normoxia, ischemia, and I/R conditions, with H₂O₂ additionally included in the viability and cTnI assays. Contractility was measured in the normoxia, ischemia, I/R, delayed normoxia, and delayed ischemia conditions.

Simulated ischemia was induced by transferring chips into a gas-tight, temperature-controlled hypoxia chamber (NOX-E.5-GTC, Noxygen, Germany) pre-equilibrated with a hypoxic gas mixture (1% O₂, 5% CO₂, 94% N₂) and replacing the culture media with glucose-free, serum-free ischemic media (DMEM, no glucose, no glutamine, no phenol red; Thermo Fisher, Waltham, MA; Cat# A1443001) to simultaneously recapitulate substrate deprivation and hypoxia (Fig S2). Successful establishment of hypoxic conditions within the chamber was confirmed by measuring dissolved oxygen levels in both the chamber atmosphere and the well media using a FOSPOR AI-300 optical oxygen-sensing probe coupled to a NeoFox-GT phase fluorimeter (Ocean Insight, Orlando, FL). Gas flow and oxygen transfer within this chamber configuration have been previously measured and validated using a calibrated computational model of the system^40^. The sealed chamber was maintained at 37°C for 2 h. For the I/R condition, ischemic media was aspirated at the end of the 2-h ischemic period, replaced with normoxic 10% FBS M199, and cultures were returned to the standard incubator for an additional 2 h of reperfusion (Fig. S2).

### Cell Viability Assay

Following each experimental condition, cell viability was assessed using a fluorescent LIVE/DEAD™ Viability/Cytotoxicity Kit (Thermo Fisher, Waltham, MA; Cat# L3224). NRVMs were incubated with 1 µM calcein-AM, 4 µM ethidium homodimer-1 (EthD-1), and 1 µg/mL Hoechst 33342 (Thermo Fisher, Waltham, MA; Cat# H1399) in warm PBS for 15 min at 37°C, protected from light. Chips were then washed once with warm PBS and immediately imaged on a fluorescence microscope (EVOS FL Imaging System). Ten randomly selected fields of view per chip were acquired. Nuclear segmentation was performed on the EthD-1 and Hoechst channels using Cellpose 2.0, an automated deep-learning-based segmentation algorithm^41,42^. Cell death was quantified from paired Hoechst (total nuclei) and EthD-1 (dead-cell marker) images. EthD-1-positive objects smaller than 25 px² were excluded as debris. Each remaining EthD-1 object was checked for pixel-level overlap with a Hoechst-positive nucleus. Total cell number was calculated as Hoechst-positive nuclei plus filtered EthD-1-positive objects, minus the number of overlapping objects, to prevent double-counting cells positive in both channels. Percent cell death was calculated as filtered EthD-1-positive objects divided by total cell number, multiplied by 100; percent viability was calculated as 100 minus percent cell death (Fig. S3).

### Cardiac Troponin I (cTnI) ELISA

Prior to fixing or live staining the cells for structural and viability analysis, 800 µL of conditioned media was collected from each experimental group at the end of the treatment period, centrifuged for 20 min at 1,000 × g (4°C) to remove cellular debris. Cardiac troponin I (cTnI) concentration was quantified using a Rat Troponin I Type 3 (Cardiac) ELISA Kit (Colorimetric; Novus Biologicals, Cat# NBP3-00484), following the manufacturer’s protocol without modification.

### Immunofluorescence and Structure Analysis

Following experimental treatments, the chips were rinsed with PBS prior to being fixed and permeabilized with warm 4% paraformaldehyde (PFA; VWR, Radnor, PA) supplemented with 0.05% Triton X-100 (Sigma-Aldrich, St. Louis, MO) for 10 min. Cells were then rinsed three times with room temperature PBS, allowing the PBS to sit for 5 min between rinses. The cells were then stained for nuclei (4′,6-diamidino-2-phenylindole dihydrochloride, DAPI; Invitrogen, Waltham, MA), actin (Alexa Fluor 488 Phalloidin; Thermo Fisher, Grand Island, NY), and sarcomeric α-actinin (Mouse Monoclonal Anti α-actinin; Sigma-Aldrich, St. Louis, MO) by incubating the coverslips in 1:200 dilutions in PBS for 1 h at room temperature. The coverslips were rinsed, and then secondary staining was performed with goat anti-mouse IgG secondary antibodies (Alexa Fluor 594; Thermo Fisher, Grand Island, NY) 1:200 dilutions in PBS for 1 h at room temperature. Coverslips were then mounted onto glass slides using ProLong Glass Antifade Mountant (Thermo Fisher, Waltham, MA) and stored at 4°C until imaging. Immunofluorescence imaging was performed on an Olympus Fluoview FV3000 confocal microscope (Olympus America, Center Valley, PA) equipped with a UPLSAPO 40× silicone-oil immersion objective, yielding a resolution of approximately 6.5 pixels/µm. Ten fields of view were randomly acquired per sample.

Sarcomeric structure was quantified using a previously described automated MATLAB image analysis pipeline^35,43,44^. Binary skeletons of α-actinin-stained z-lines were extracted from each image and analyzed to compute five metrics^38,39^. Z-line density per nucleus was calculated as the number of z-line pixels divided by the total number of nuclei in the same field of view, providing an index of sarcomeric content normalized to nuclei number. Z-line orientational order parameter (OOP) quantified the degree of z-line alignment. Z-line fraction quantified the proportion of an α-actinin signal residing in well-formed z-lines. Z-line integrity captured the extent of z-line breakdown and the accumulation of diffuse α-actinin signal. Mean continuous z-line length quantified the average length of uninterrupted z-line segments. Each metric was averaged across the ten fields of view acquired per chip (Fig. S1).

Cardiomyocyte area was quantified from the imaging dataset using a decision tree classifier implemented in a custom MATLAB script, following a previously established approach^39^. Approximately 1% of the dataset was manually annotated to label cardiomyocyte-containing regions, and these annotations were used to train and optimize the classifier, which achieved a classification accuracy of 98%. The trained classifier was then applied across the full dataset to identify cardiomyocyte regions and quantify the number of actin-positive pixels contained within them.

### MTF Contractility Measurements

Contractile stress was measured using the MTF platform immediately before and immediately after each experimental condition, enabling within-substrate paired comparisons. Prior to imaging, the PIPAAm sacrificial layer was dissolved by briefly immersing the coverslip in room-temperature PBS (<25°C), causing the PDMS film to delaminate from the underlying glass. Individual MTF cantilevers were then excised with a razor blade and transferred into 35 mm Petri dishes containing the appropriate culture media for the corresponding experimental condition. Petri dishes were placed on a temperature-controlled heating stage positioned beneath a stereo microscope (Olympus America, Center Valley, PA). A mounted camera (Basler, Germany) recorded film deflection from above the microscope at 100 frames per second.

Tissues were field stimulated using a pulsatile field stimulator (Myopacer, IonOptix, Westwood, MA) delivered through two carbon electrodes (McMaster-Carr, Douglasville, GA) positioned 1.5 cm apart. Contractile behavior was recorded while films were externally paced at 2 Hz with a stimulation voltage of 20 V. Sarcomeric stress was subsequently calculated from the recorded video data using a previously described automated analysis pipeline implemented in ImageJ and MATLAB^35^ (Fig. S1).

Average sarcomeric force was derived by combining the contractility data (monolayer-level stress) with the structural data on cardiomyocyte-occupied area obtained from the classification analysis described above^37,39^. Cardiomyocyte area (*A_CM_*) was defined as the fraction of each field of view occupied by actin-positive pixels classified as cardiomyocytes, excluding fibronectin-only and non-cardiomyocyte regions (Fig. S4). Cardiomyocyte area was averaged across the fields of view acquired per chip. Sarcomeric stress was then normalized to this fraction to yield average sarcomeric stress, as previously described^39^.

### Statistical Analysis

All data are expressed as mean ± standard deviation (SD) unless otherwise noted. Statistical comparisons across conditions were performed using one-way and two-way analysis of variance (ANOVA) followed by Tukey’s post hoc test for multiple comparisons. Statistical significance was set at α = 0.05 and is indicated as: *p < 0.05, **p < 0.01, ***p < 0.001, ****p < 0.0001. All analyses were conducted in GraphPad Prism (version 10). The number of independent biological replicates (n) and the unit of replication for each assay are specified in the corresponding figure legends.

## 3. RESULTS

### A. Ischemic sarcomere loss occurs without architectural disorganization

To characterize the structural consequences of ischemic injury, NRVMs cultured on MTFs were subjected to 2 hours of ischemia or 2 hours of ischemia followed by 2 hours of reperfusion (I/R), and subsequently immunostained for sarcomeric α-actinin to visualize z-line organization (Fig. 1A). Z-line density per nucleus was significantly reduced under both ischemic and I/R conditions relative to normoxia. The magnitude of this deficit was comparable between the two injured groups, indicating that reperfusion neither exacerbated nor reversed the ischemic loss (Fig. 1B). In contrast, metrics reflecting the regularity and integrity of residual sarcomeric architecture were unaffected by injury (Fig. 1C–E) as would be expected for only 2 hours hypoxia^38^. Indeed, the z-line orientational order parameter (OOP), which quantifies the degree of z-line alignment, was comparable across all three conditions (Fig. 1C), indicating that the sarcomeres remaining after ischemia and reperfusion retain their organized registry rather than becoming structurally disordered. Similarly, z-line fraction, which represents the proportion of total α-actinin signal residing in well-formed z-lines, was maintained (Fig. 1D), demonstrating that the surviving sarcomeric network remains well matured and does not undergo fragmentation. Mean continuous z-line length was likewise unchanged across conditions (Fig. S5), indicating that surviving z-line retained their longitudinal continuity. The same held for z-line integrity, which captures the extent of z-line breakdown and the accumulation of diffuse α-actinin signal (Fig. 1E), arguing against dissolution or progressive degradation. Together, these findings establish that the observed reduction in sarcomeric density reflects a net loss of cardiomyocytes rather than structural degradation of sarcomeres within cardiomyocyte.

**Figure 1.**
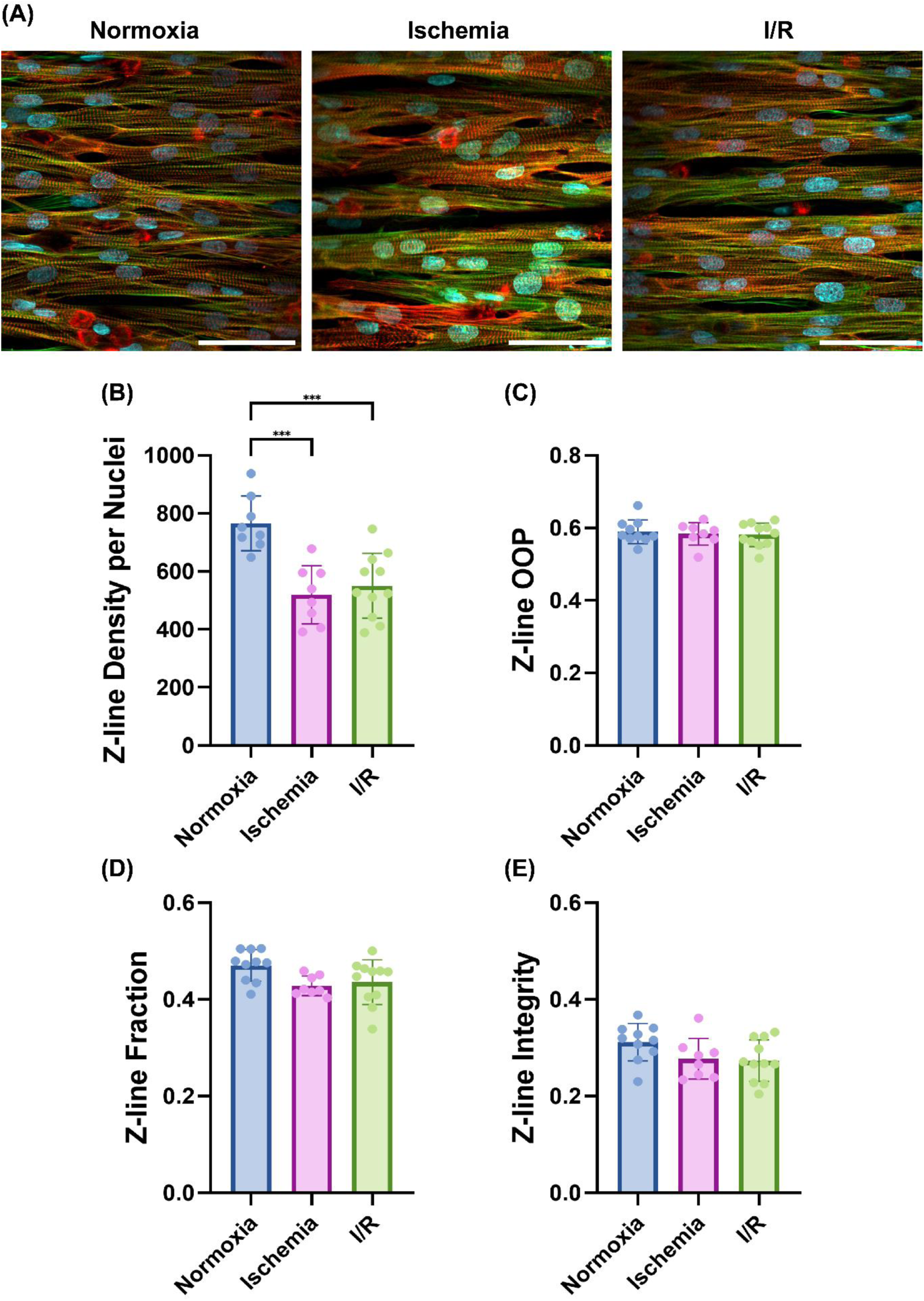
Sarcomeric z-line structural analysis. (A) Representative immunofluorescent images of NRVMs under normoxia, ischemia, and I/R conditions; stained for nuclei (blue), actin (green), and α-actinin (red). Scale bar = 50 μm. (B) Z-line density per nuclei. (C) Z-line orientational order parameter (OOP). (D) Z-line fraction. (E) Z-line integrity. Statistical significance was determined by one-way ANOVA with Tukey’s post hoc multiple comparison tests. ***p<0.001. n = 8-10 coverslips; 4 harvests per group

### B. Ischemia induces cell death and cTnI release, both exacerbated by reperfusion

To assess the degree of cardiomyocyte injury across conditions, cell viability and conditioned media cardiac troponin I (cTnI) concentration were quantified as complementary indices of membrane integrity and myofilament protein release, respectively (Fig. 2). LIVE/DEAD fluorescence staining revealed a condition-dependent loss of viability across all injury groups relative to normoxia, with H₂O₂ treatment producing the most severe cell death, followed by I/R, then ischemia (Fig. 2B). Viability in I/R was significantly lower than in ischemia, indicating that reperfusion drives further cell death beyond that sustained during ischemia alone.

**Figure 2.**
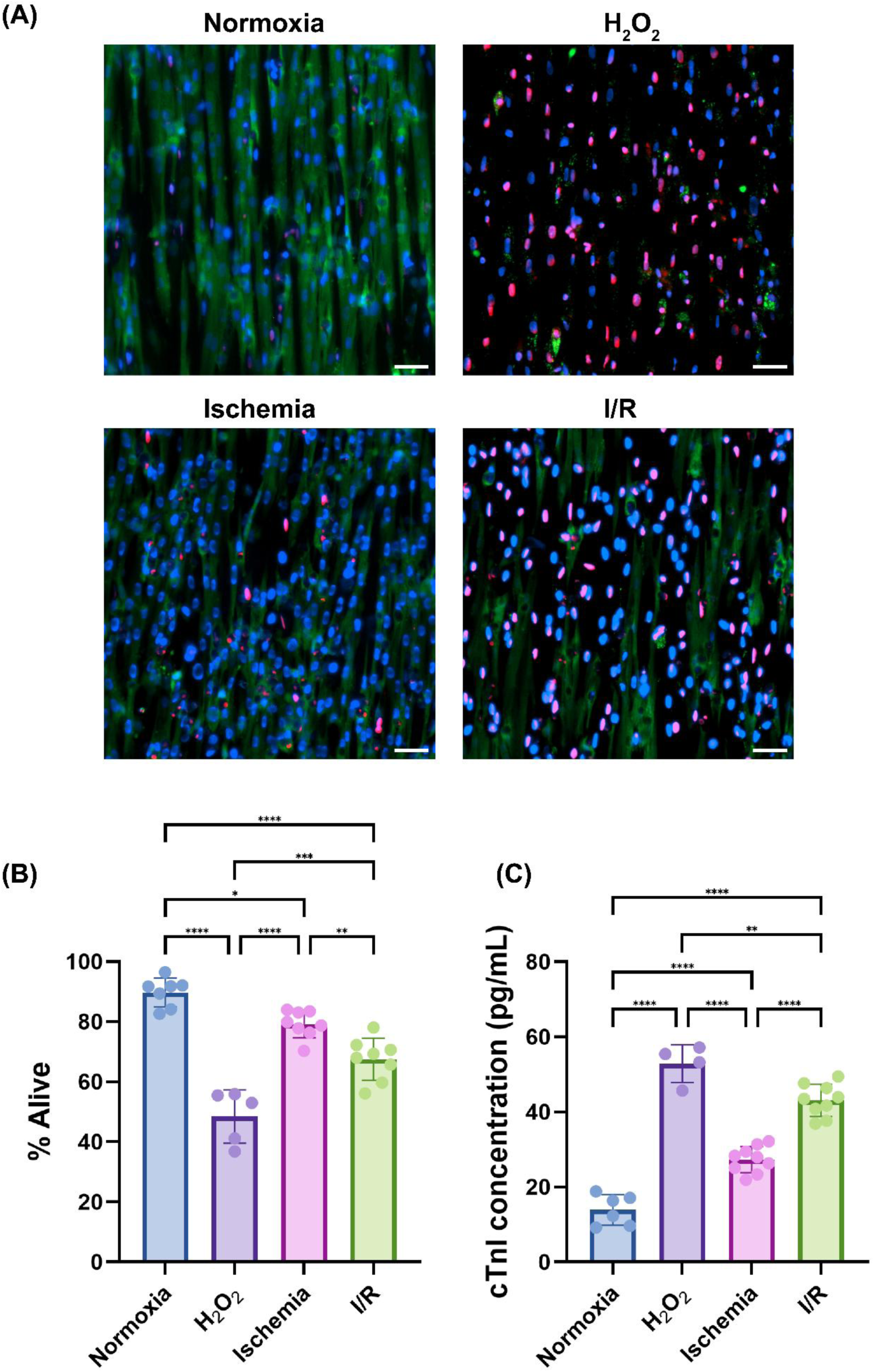
Cell viability and cTnI injury biomarker analysis. (A) Representative live/dead fluorescent images of NRVMs under normoxia, H₂O₂, ischemia, and I/R conditions; stained for Hoechst (total nuclei, blue), calcein-AM (live cells, green), and ethidium homodimer-1 (EthD-1; dead cells, red). Scale bar = 100 μm. (B) Percentage of viable cells across conditions, quantified from Hoechst and EthD-1 channels. (C) Conditioned media cTnI concentration (pg/mL) across conditions. Statistical significance was determined by one-way ANOVA with Tukey’s post hoc multiple comparison test. *p < 0.05, **p < 0.01, ***p<0.001, ****p < 0.0001. *n* = 5-8 coverslips; 4 harvests per group

Conditioned media cTnI concentration mirrored this pattern of severity in a graded and concordant manner (Fig. 2C). Normoxia exhibited the lowest cTnI release, with successively higher in ischemia, I/R, and H₂O₂. The step-up in cTnI from ischemia to I/R was also statistically significant, confirming that reperfusion drives additional myofilament protein release beyond what ischemia alone produces. Conditioned media cTnI therefore represents injury severity with the same resolution as direct viability assessment, validating a clinically established biomarker assay as a non-destructive readout in this heart-on-a-chip NAM.

### C. Reperfusion dissociates systolic and diastolic stress and destabilizes beating

Contractile function was assessed using the cardiac muscular thin film (MTF) platform, which enables measurement of cardiac stress before and after experimental conditions in the same preparation (Fig. 3A). Electrical field stimulation at 2 Hz induced contraction of the MTFs, causing upward curling away from the coverslip that permitted measurement of systolic and diastolic stresses. Stress ratios (σ_after_/σ_before_) were computed for each film to normalize treatment-attributable changes to each film’s own pre-condition baseline, controlling for inter-film variability. Because NRVMs continue to mature in culture over time, a stress ratio change observed between baseline and endpoint could in principle reflect time-dependent changes in the preparation itself rather than a genuine effect of ischemia or reperfusion. The delayed normoxia and delayed ischemia groups control this possibility. Both begin at t₂, the exact timepoint at which I/R reperfusion begins and are measured at t₄ using t₂ as their condition-specific baseline, equalizing time-in-culture across groups and isolating the reperfusion stimulus from cumulative maturation effects (Fig. 3B).

**Figure 3.**
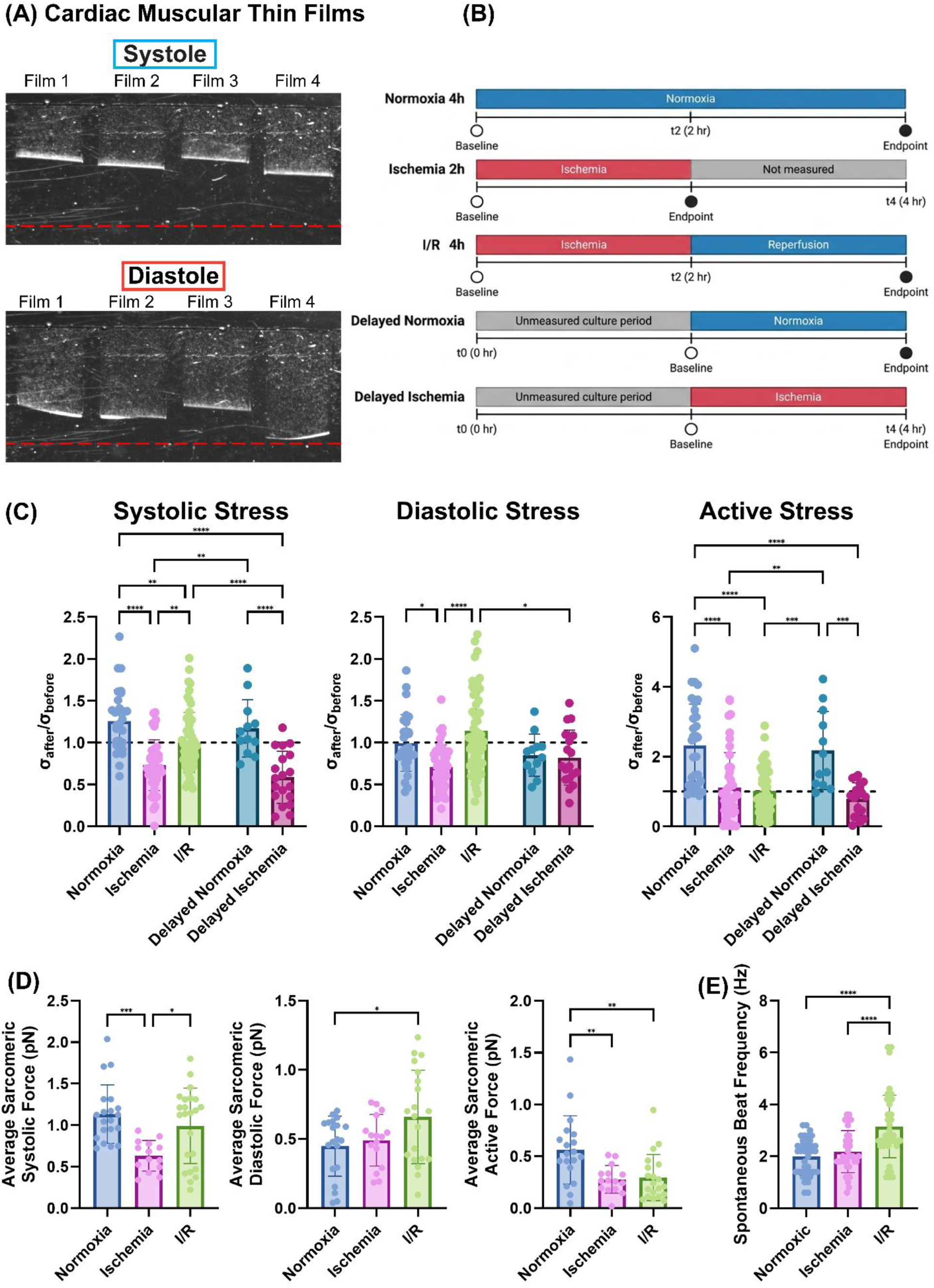
Contractile function analysis of NRVMs on the MTF platform. (A) Representative brightfield images of MTFs at systole and diastole. (B) Schematic of the experimental timeline for each condition (Normoxia 4h, Ischemia 2h, I/R 4h, Delayed Normoxia, Delayed Ischemia), indicating the time points at which baseline and endpoint stress measurements were acquired. (C) Normalized stress ratios (σ_after_/σ_before_) for systolic, diastolic, and active stress across conditions; dashed line at 1.0 indicates no change from baseline. (D) Average sarcomeric force (pN) for systolic, diastolic, and active stress across conditions. (E) Beat frequency (Hz) before and after intervention across conditions. Statistical significance was determined by one-way and two-way ANOVA with Tukey’s post hoc multiple comparison test. *p < 0.05, **p < 0.01, ***p<0.001, ****p < 0.0001. *n* = 30-50 films; 3-5 harvests per group

Systolic stress ratio was significantly reduced in ischemia relative to normoxia, reflecting ischemia-induced depression of the contractile apparatus (Fig. 3C). Following reperfusion, the systolic ratio in I/R was significantly greater than in ischemia, indicating partial recovery, but remained significantly below normoxia, indicating incomplete functional restitution. I/R also differed significantly from delayed ischemia, supporting that this partial recovery reflects a genuine reperfusion effect rather than time-in-culture maturation of the NRVMs. Average sarcomeric systolic force mirrored this pattern, with ischemia exhibiting the lowest values and I/R showing partial recovery (Fig. 3D).

Diastolic stress ratio was significantly reduced in ischemia relative to normoxia but increased significantly upon reperfusion in I/R relative to ischemia, with values trending above the normoxic baseline, although the direct Normoxia–I/R comparison did not reach significance (Fig. 3C). I/R also differed significantly from delayed ischemia, indicating that this diastolic elevation is specific to reperfusion rather than elapsed culture time. Ischemia alone did not elevate diastolic stress. It was reduced relative to normoxia (Fig. 3C) and unchanged at the level of individual sarcomere (Fig. 3D). Following reperfusion, however, diastolic force per sarcomere increased above normoxic levels (Fig. 3D). This selective elevation of diastolic stress following reperfusion, without a comparable change during ischemia alone, identifies reperfusion as a primary driver of impaired myocardial relaxation in this model.

Active stress ratio, which represents the difference between systolic and diastolic stress normalized to baseline and reflects net contractile reserve, was markedly suppressed in both ischemia and I/R relative to normoxia, with no significant difference between the two injury conditions (Fig. 3C). I/R also differed significantly from delayed normoxia, indicating that this deficit cannot be attributed to time in culture. Average active sarcomeric force was similarly and significantly reduced in both ischemia and I/R relative to normoxia (Fig. 3D). Because active stress reflects the difference between systolic and diastolic stress, the partial systolic recovery in I/R was offset by a concurrent rise in diastolic stress, leaving active stress suppressed relative to normoxia. Systolic indices alone would overestimate functional recovery after reperfusion. In the absence of electrical stimulation, spontaneous beat frequency did not differ between normoxia and ischemia (Fig. 3E). In contrast, spontaneous frequency after I/R was significantly higher than both normoxia and ischemia. The post-I/R group also exhibited markedly increased variability in beat frequency, suggesting heterogeneous electrophysiological responses across preparations. Together, these findings demonstrate that ischemia and reperfusion produce mechanistically distinct contractile phenotypes. Ischemia primarily suppresses systolic contractile output, whereas reperfusion additionally impairs diastolic relaxation and alters beating rate, while active stress remains suppressed under both conditions. Each of these features recapitulates a clinically recognized aspect of myocardial ischemia-reperfusion injury.

## 4. DISCUSSION

This study demonstrates that a single heart-on-a-chip NAM can resolve the structural, biochemical, and contractile dimensions of myocardial I/R injury. Rather than reporting a single index of injury, the platform decomposes dysfunction into components that are otherwise entangled in the intact heart, each mapping onto a distinct clinical phenotype. Ischemia and reperfusion therefore produced qualitatively distinct injury states: under ischemia, all readouts moved together, consistent with cell loss, whereas after reperfusion they diverged, so that no single readout could capture the full injury. Many of these findings align with existing *in vivo* and clinical observations, supporting the platform’s translational relevance. The work, therefore, contributes an experimental framework in which mechanistically distinct forms of cardiac dysfunction can be captured, distinguished, and studied within a shared preparation.

Ischemia in the absence of reperfusion is the pathophysiology of Type 2 MI^45,46^, and the present data reproduced it across biochemical, structural, and contractile readouts. Simulated ischemia reduced cell viability relative to normoxia (Fig. 2B) and conditioned media cTnI increased in parallel (Fig. 2C), together representing the *in vitro* analog of ischemic injury burden accrued before flow is restored^9^. Notably, sarcolemmal rupture and myofilament protein release tracked together confirming the protocol recapitulated both processes simultaneously, as occurs in ischemic myocardium *in vivo*^47–49^. The structural data then specified where this injury did not occur. Z-line density per nucleus declined significantly under ischemia (Fig. 1B), while z-line OOP, z-line fraction, and z-line integrity remained stable (Fig. 1C–E). Because z-line density is nuclei-normalized, its reduction reflected net loss of sarcomere-containing cells rather than sarcomeric disassembly within survivors, a pattern consistent with the heterogeneous, patchy myocyte loss that characterizes the evolving ischemic wavefront in the myocardium^50,51^. Functionally, systolic, diastolic, and active stress ratios were each significantly reduced under ischemia (Fig. 3C), with average systolic and active force declining in parallel (Fig. 3D), while spontaneous beat frequency remained unchanged (Fig. 3E). This pattern is consistent with tissue containing fewer viable sarcomere-containing cells, generating proportionately less total output while continuing to beat at a stable rate. Reduced contraction is also the earliest detectable consequence of coronary occlusion in patients, with regional hypokinesis appearing on echocardiography within seconds of balloon inflation and preceding both electrocardiographic change and angina^52,53^. Taken together, the structural, biochemical, and contractile readouts under ischemia all pointed to a single explanation, loss of viable cardiomyocytes, with rhythm preserved.

The I/R condition was not a more severe version of ischemia but a qualitatively different pathological state. Viability declined significantly further from ischemia to I/R (Fig. 2B), reflecting an additional wave of cardiomyocyte death, believed to be driven by mPTP opening and ROS generation at the moment of pH normalization upon reperfusion^8,9,54,55^, with cTnI release increase in parallel (Fig. 2C). Critically, this incremental rise above the ischemic level, rather than any absolute magnitude of cell death that maps directly onto the Type 4a periprocedural MI framework of the Fourth Universal Definition of MI^45,56^, which is the diagnostic category for reperfusion-associated myonecrosis following percutaneous coronary intervention that carries prognostic significance independent of the presenting infarct^56–60^. Structurally, however, nothing changed beyond the extra cell death. Z-line density remained reduced without differing appreciably from ischemia (Fig. 1B), and the architectural metrics were stable across all conditions (Fig. 1C–E). Thus, reperfusion neither restored the sarcomere-containing population lost during ischemia nor degraded the architecture of surviving cells, so every functional change observed in I/R occurred on a numerically depleted but architecturally intact myocyte population. Importantly, every functional change observed in I/R thus occurred in a depleted but architecturally intact cardiomyocyte population, indicating that the subsequent dysfunction was not attributable to sarcomere disorganization, a conclusion the contractile data alone could not support.

Against this backdrop, systolic stress ratio recovered partially in I/R relative to ischemia (Fig. 3C), with absolute sarcomeric force recovering correspondingly (Fig. 3D), indicating that surviving, structurally ordered cells regained partial contractile output upon restoration of flow. This incomplete recovery on an intact sarcomeric substrate is the defining signature of myocardial stunning^61–64^. The residual deficit has been attributed to myofilament-intrinsic contractile depression whose mechanistic basis remains debated, whether a reduction in maximum Ca²⁺-activated force at preserved Ca²⁺ sensitivity^65^ or concurrent reductions in both Fmax and myofilament Ca²⁺ sensitivity^66–68^. Common to both accounts is a disturbance in calcium handling at reperfusion, and the diastolic stress data showed its consequences extended beyond the systolic deficit.

Diastolic stress ratio reversed direction with reperfusion, falling below normoxia under ischemia but increasing significantly above the ischemic level and trending above baseline in I/R (Fig. 3C), with absolute diastolic force elevated relative to normoxia (Fig. 3D). Excess cytosolic Ca²⁺ at reperfusion overwhelming SERCA2a-mediated re-sequestration would sustain crossbridge engagement into diastole on architecturally normal sarcomeres, producing the elevation in diastolic stress observed here^67,69–71^. Clinically, this maps onto the restrictive left ventricular filling pattern identified by the MeRGE AMI Collaborators (2008)^72^ as an independent predictor of post-MI mortality and heart failure, and connects mechanistically to the diastolic dysfunction at the core of heart failure with preserved ejection fraction, whose post-MI incidence remains substantial despite successful revascularization^6,73,74^. The same disturbance accelerated spontaneous rhythm. Beat frequency elevated significantly in I/R with markedly increased inter-preparation variability, while remaining stable across normoxia and ischemia (Fig. 3E). Cytosolic Ca²⁺ overload is believed to trigger spontaneous sarcoplasmic reticulum calcium release during diastole, generating delayed after depolarizations via Na⁺/Ca²⁺ exchanger reversal and triggered activity, the dominant mechanism of clinical reperfusion arrhythmias and mechanistically distinct from the reentrant arrhythmias of ischemia^75–79^. The diastolic elevation and increased beat frequency emerged together and only in I/R is, therefore, unlikely to be coincidental, since both are downstream of a single calcium-handling disturbance, and neither would be mechanistically interpretable without the other.

Active stress captured the consequence of systolic and diastolic stress moving in opposite directions. Active stress ratio was persistently suppressed in I/R despite partial systolic recovery (Fig. 3C), because the concurrent diastolic overshoot arithmetically offset systolic gains. Active stress in this platform is a cell-level index of contractile reserve rather than an ejection fraction surrogate, since the MTF captures no volumetric displacement, though comparable dissociations between systolic indices and net output carry prognostic weight clinically^80^. A systolic-only readout would have reported this condition as substantially recovered, whereas the persistent active stress deficit shows that apparent systolic recovery coexisted with a genuine loss of net contractile output. Consequently, that deficit was visible only because systolic and diastolic stress were resolved within a single heart-on-a-chip, a distinction a platform built around one global index could not have made.

## 5. CONCLUSION

Collectively, this work establishes the heart-on-a-chip platform as a NAM in which the structural, biochemical, and contractile dimensions of myocardial ischemia and reperfusion injury can be resolved within a single preparation. By separating readouts that remain entangled in the intact heart, the platform makes it possible to distinguish forms of contractile dysfunction that arise from mechanistically distinct origins yet present similarly at the level of global function. This resolution matters for therapeutic screening, since an intervention that improves one dimension of function while worsening another cannot be identified by a platform that reports only one, and cardioprotective strategies have repeatedly succeeded against single experimental preclinical endpoints without translating to patient benefit^14–18^. The optical accessibility of the MTF format further supports ratio-metric calcium imaging alongside contractile measurement, making direct measurement of calcium handling the immediate next development. Building on this foundation, the platform is positioned for pharmacological validation, testing candidate cardioprotective agents against the full injury phenotype, and determining whether apparent efficacy on one readout persists or is offset across the others. The value of the heart-on-a-chip NAM therefore lies not only in what it reveals about ischemia and reperfusion as distinct pathological states, but in the standard it sets for how future cardioprotective interventions should be evaluated.

## ACKNOWLEDGEMENTS

The authors thank Nida T. Qayyum for characterizing gas flow and oxygen transport within the hypoxia chamber used in this study.

